# Seeing new ‘stuff’ for the very first time

**DOI:** 10.64898/2026.11.19.767518

**Authors:** Jacob R. Cheeseman, Filipp Schmidt, Hua-Chun Sun, Chenxi Liao, Bei Xiao, Roland W. Fleming

## Abstract

Humans are excellent at recognizing materials^1^ like food, soil or paper, which is crucial for many everyday tasks^2^. We generally enjoy vivid intuitions about what surfaces would feel like, without reaching out and touching them. While this capability is understandable for familiar materials, how does the brain interpret stuff we have never encountered before? We hypothesized that the need to distinguish useful or hazardous substances by sight led to the ability to infer material properties from the very first encounter. To test this, we used generative AI to synthesize realistic images of new ‘hybrid’ materials, by fusing characteristics of images from different ‘parent’ categories (e.g., bark × carpet). Even though the hybrid materials have never existed in the real world—and observers judged them to be far less familiar than their parents—we found that they elicited a rich and compelling phenomenology, and yielded consistent inferences about their likely physical and multisensory properties, such as what they would feel like to touch. Their perceived properties occupied the same broad multidimensional space as familiar materials, whereas phase-scrambled controls did not. Our findings suggest that the visual system exploits generalizable relationships between visual appearance and physical properties, enabling instantaneous interpretation of new kinds of ‘stuff.’

## Introduction

Recognizing materials and their physical characteristics is one of our most important visual capabilities, essential for food selection, tool use and many other everyday behaviours^3^. Previous studies have extensively mapped out how the human visual system represents familiar material categories, like paper, grass and water, in a multidimensional ‘material appearance space‘^2,4–7^. The representational relationships between materials—which underlie both similarity judgments and the recognition of familiar materials—presumably emerged through extensive multisensory experience over both evolution and our individual lifetimes. Yet what happens when we encounter a completely new material for the first time? Throughout our lives, we frequently encounter unfamiliar foodstuffs or textiles, and indeed, in the modern world new materials emerge surprisingly often (e.g., holograms, boba tea, transparent wood). Can we work out their properties, such as how they would feel, without even touching them? If material perception depended solely on recognizing familiar categories and retrieving the learned properties from memory, genuinely novel materials would leave us with little to go on.

One might imagine that learning a new material requires active, multisensory exploration like prodding, stroking or tilting it in the light^8^. Indeed, playing with materials, from wooden blocks to food to playdough, is a preferred activity of young children, presumably to form or fine-tune sophisticated internal models^9^. Yet, we hypothesized that adults’ extensive experience with materials may enable us to recognize new ‘stuff’ without active exploration. Specifically, we might be able to make sense of novel materials through compositional generalisation^10^, i.e., recombining the features of familiar materials. Thus, understanding how we perceive novel materials could not only provide insights into the representation of unpopulated regions of material space, but also hint at how we achieve ‘zero-shot’ generalization more broadly.

Until recently, creating photorealistic novel materials would be technically infeasible. To overcome this, we prompted a generative AI to synthesize images by combining visual characteristics of familiar material categories (‘parents’) in ways that do not ordinarily occur in the world. We first tested whether the resulting ‘hybrid’ materials appeared genuinely unfamiliar. We then asked the more critical question: despite their novelty, could observers form vivid and systematic impressions of their physical and multisensory properties?

## Methods and Results

To create novel but visually coherent materials, we used Midjourney^11^ to synthesize photorealistic images of 26 familiar material categories (‘parent’ images), such as wood, paper, paint, skin, and ceramics. We then used Midjourney to blend pairs of parent images to create ‘hybrid’ images (see **Figure 1** for examples). For each parent and hybrid class, we generated four similar but distinct examples. We manually excluded images that appeared as a patchwork of distinct materials rather than a single uniform surface, leaving a total of 369 hybrids (all stimuli, data, text prompts and code available here: https://doi.org/10.5281/zenodo.23395585).

**Figure 1.**
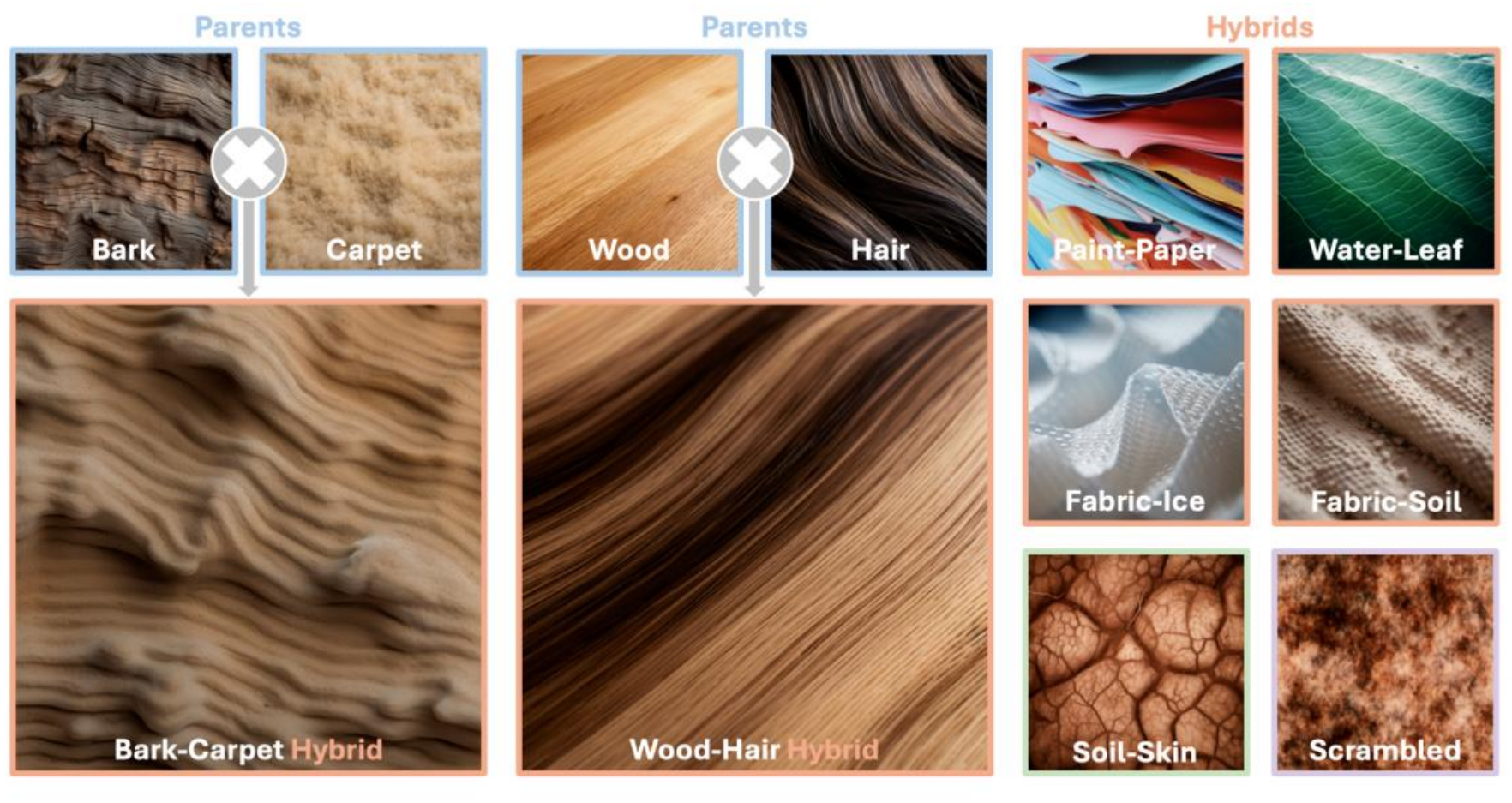
Example AI-generated ‘parent’ and ‘hybrid’ materials. A phase-scrambled control image is shown for ‘Soil-Skin’ (bottom right). Note the vivid impression of coherent, photorealistic surfaces with recognizable properties in both familiar (parent) and novel (hybrid) materials, but not control).

We first tested whether the hybrid materials appeared truly novel. We would expect two signatures of novelty: observers should (1) judge them to be less familiar than parent materials, and (2) have greater difficulty describing them consistently. To test this, we showed twenty participants 2250 unique pairs of images (one parent and one unrelated hybrid per trial), and asked them to report which appeared more familiar (see **SI** for details). A separate group of forty participants provided brief text descriptions for a randomly-selected subset of 138 images.

The results (**Figure 2A**) show that hybrid images were less familiar than parent images (paired *t*-test: *t*(19) = −19.61, *p* < .001) and took on average 3.5 seconds longer to describe (paired *t*-test: *t*(9) = 2.62, *p* < 0.05). Descriptions of hybrids were also less consistent than descriptions of parent images (paired *t*-tests: *t*(9) = −6.10, *p* < 0.001), measured by the cosine similarity between sentence embeddings of participants’ descriptions (**Figure 2B**)^12^. This was presumably because it was harder to work out how to describe these unusual materials. Yet, descriptions of both hybrid and parent images were significantly more consistent than random samples of the responses (paired *t*-test for parent: *t*(9) = 19.13, *p* < 0.001; and for hybrid: *t*(9) = 12.31, *p* < 0.001). Descriptions of hybrid materials were also more similar to their parents, than parents were to one another (paired t-test: *t*(7) = 4.16, *p* < 0.01). Together these findings suggest that hybridization preserved recognizable characteristics of the parent materials while combining them into unfamiliar configurations. The stimuli therefore provide a crucial test case: materials that observers could not readily identify, but whose appearance nevertheless contained interpretable features inherited from familiar materials.

**Figure 2.**
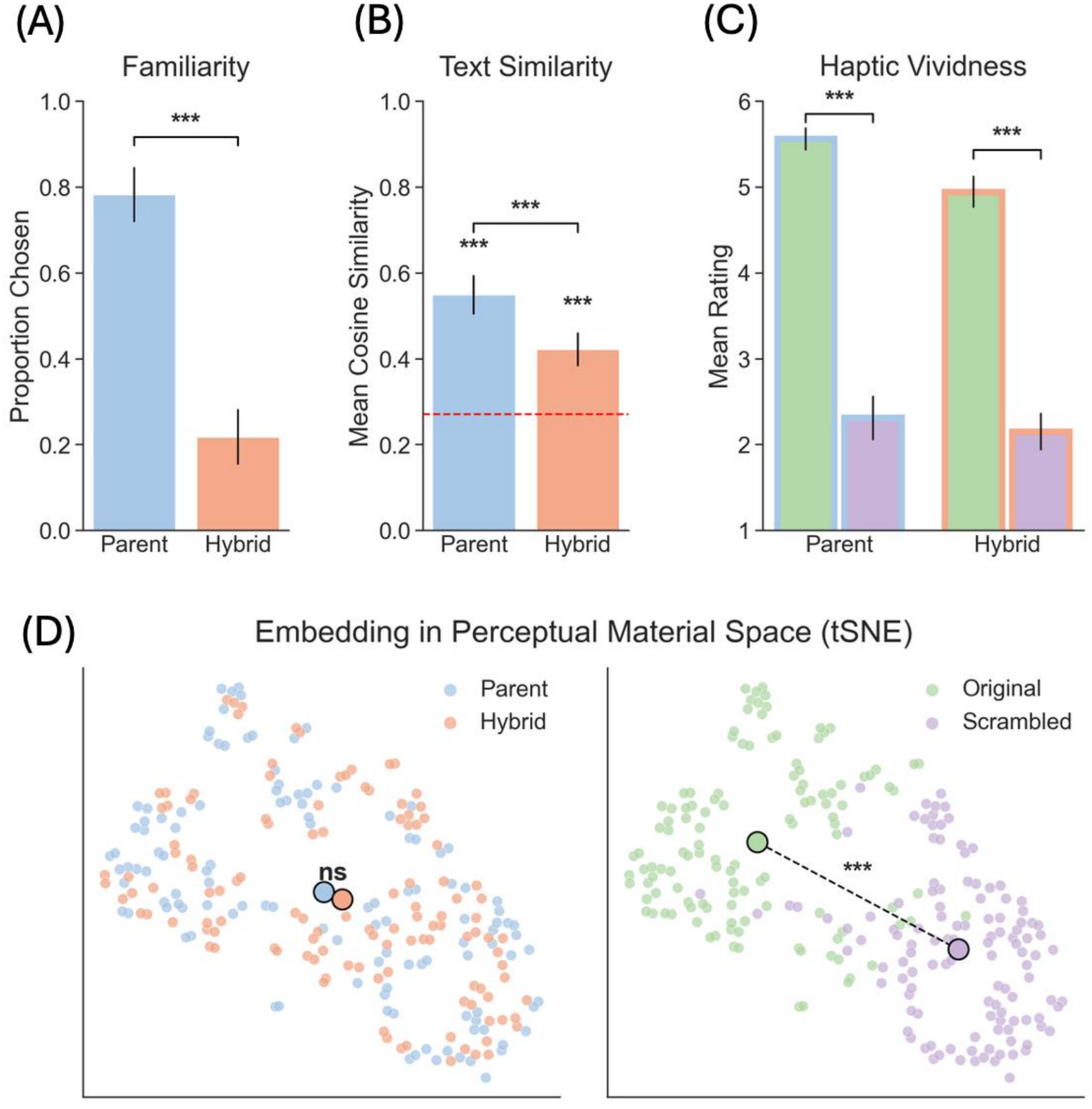
Experimental results. **(A)** Familiarity ratings were substantially higher for parents than hybrids. **(B)** Text descriptions of parents were significantly more similar than hybrids, and both were described more consistently than chance (red dashed line). **(C)** Compared with phase-scrambled scrambled images, the original versions evoked a significantly more vivid impression of what they would feel like to touch. **(D)** When perceptual ratings of images are visualized in latent space (using tSNE), parents and hybrids overlap, while ratings of original and scrambled images form separated clusters.

Could observers use these image features to infer properties of materials they had never encountered? To test this, we asked twenty more participants to rate nine physical and multisensory characteristics of both parent and hybrid images (photorealistic/pixelated; 3D relief/flat; soft/hard; rough/smooth; solid/crumbly, grainy; warm/cold; wet/dry; appealing/repulsive to touch; vividness/vagueness to touch; see **SI** for details). As a control, we included phase-scrambled versions of the images (see “**Scrambled**” inset, **Figure 1**). These had the same color distribution and amplitude spectrum as the source images, but lacked coherent spatial organization, and were rated as significantly less 3D (paired *t*-test: *t*(119) = −23.66, *p* < 0.001). This allowed us to ask whether vivid material impressions depended on coherent surface appearance or merely low-level image statistics.

Despite never having seen or touched the hybrid materials, observers formed vivid impressions of how they would feel (**Figure 2C**). Ratings of the hybrid materials were similar to those elicited by familiar parent materials and significantly higher than scrambled controls (paired *t*-test: *t*(119) = 7.78, *p* < 0.001). Thus, even unfamiliar materials could evoke compelling expectations about tactile properties.

Finally, we asked whether novel and familiar materials elicit fundamentally different kinds of percepts, or whether both are represented within a common perceptual space. We represented each stimulus by its ratings along the nine perceptual dimensions (**Figure 2D**). Overall, we found that hybrid and parent images occupied broadly overlapping regions in this ‘perceptual material space’ (little to no evidence for a difference, BF_10_ = 0.53; classification accuracy = 56.5%). In contrast, the phase-scrambled versions of the images were readily separable from the originals (extreme evidence for a difference, BF_10_ = 1.58 × 10^48^; classification accuracy = 94.6%). Thus, novelty did not place the hybrids outside the range of ordinary materials. Instead, unfamiliar materials were accommodated within much the same space of perceived physical and multisensory properties as familiar ones.

## Discussion

Our results show observers can form rich and systematic impressions of materials that neither they nor any evolutionary ancestor has encountered before. While familiar materials can be recalled from previous experiences, this does not work with novel materials. Instead, the visual system must generalize from familiar materials to novel combinations of appearance characteristics. On this account, experience with materials such as wood, skin, ceramic or moss does more than establish fixed associations between particular categories and their multisensory properties. It also establishes a structured mapping between visual appearance and likely physical and multisensory properties, allowing a novel material to be interpreted according to where its appearance falls within this learned structure (like a continuous manifold)—even when the material itself cannot be recognized.

This hints at a form of compositional generalization, in which familiar appearance characteristics can be recombined in unfamiliar ways^13^. More broadly, compositional representations could enable generalizing outside the distribution of previous experiences, allowing us to imagine and reason about not just novel materials but also creatures, objects, places or situations. Generative AI models are powerful tools for probing this ability by creating forms of visual novelty that are difficult to manufacture in the real world, such as the hybrids of textile, soil, snow or ceramic that we used here. Such approaches could also be used to investigate zero-shot inferences and out-of-distribution generalization in other domains.

## Acknowledgments

This work was funded by the Deutsche Forschungsgemeinschaft (German Research Foundation, DFG) under Germany’s Excellence Strategy (EXC 3066/1 “The Adaptive Mind”, Project No. 533717223) and the Research Training Group (RTG 3147 “PIMON”, Project No. 547865311), as well as The European Research Council (ERC) Advanced Grant “STUFF” (project number ERC-2022-AdG-101098225), the European Union’s Horizon Europe research and innovation programme under the Marie Skłodowska-Curie grant agreement No. 101226908 (‘EXPLORA’), and by the National Institutes of Health grant 1R15EY033512.

