## Supplemental Information for "Seeing new ‘stuff’ for the very first time"

### Stimuli

Stimuli were generated by prompting Midjourney v5.2 [1] with text prompts specifying closeup photographs of 26 categories of familiar materials; these categories have been featured in previous studies of visual material perception [2]. Example images of each material class are shown in **Figure S1**, and the text prompts used to generate these are listed in **Table S1**. By default, Midjourney outputs four slightly different variant images for each text input; these 'parent' images* were then used to create 'hybrid' images using Midjourney's built-in 'blend' function, which takes two images as input, and outputs four slightly different images that combine features of the input images. The 'blend' function assigns roughly equal weight to input images by default, meaning that in principle, the two 'parents' contributed equally to each 'hybrid' combination. Hybrid images were manually reviewed by the research team in order to exclude images that depict two or more distinct materials, rather than a single uniform 'hybrid' material. After this process, the filtered image set consisted of four variants of 26 parent classes and 369 hybrids, or 1580 images in total. Phase-scrambled and color-histogram-matched versions of a subset of these images were also created for Experiment 3. At the start of each experiment participants were instructed to place a credit card (or another card of equivalent size) on their display screen, and adjust the length and width of a rectangle to match the size of the card. This measurement was used to calibrate the size of the images such that they were approximately the same size for different displays (see https://pavlovia.org/Wake/screenscale).

*Parent images are referred to as 'base' images throughout the code and data files.

**Procedure**

Participants were recruited online using Prolific (prolific.co) and were paid €12 per hour; they were required to have native fluency in English and a desktop or laptop computer. All experimental procedures were approved by the Justus Liebig University Giessen Psychology Department Ethics Board and conformed with the guidelines of the American Psychological Association (Version 2017) and the Declaration of Helsinki (Version 2013, excluding pre-registration). Informed consent was obtained from all participants.

*Experiment 1 (Familiarity 2AFC task)*: The filtered image set was randomly sampled to yield 10 disjoint batches of 225 parent-hybrid image pairs. Each batch was presented twice to two participants. In total, 20 adults (8 female, age $M$ = 37.45, $SD$ = 11.72) were presented with two images on each trial: one 'parent' and one 'hybrid', selected at random from different categories (i.e., the parent was not the parent of the hybrid it was paired with). Their task was to indicate which of the two images depicted a material that was more familiar to them, by pressing a corresponding key.

*Experiment 2 (Free-text description task)*: The filtered image set was randomly sampled to yield 10 batches of 25 images (12 parent and 13 hybrid images). Parent images were sampled with replacement across batches; hybrids were not. Each batch was presented once to four participants. In total, 40 adults (11 female, age $M$ = 31.90, $SD$ = 8.57) were presented with a single image on each trial, either a 'parent' or a 'hybrid' image selected at random. Participants were instructed to "briefly describe the material shown in the image", and could type their responses into a text box below the image. Images were displayed for up to 1 min, after which they could continue writing their description for an unlimited duration. Participants were allowed to enter their response and proceed to the next trial before 1 min had elapsed.

*Experiment 3 (9-dimension rating task)*: The filtered image set was randomly sampled to yield 10 disjoint batches of 24 images, each of which consisted of 6 parents (2 variants of 3 categories), 6 hybrids (2 variants of 3 categories), and phase-scrambled versions of these images. A separate batch of 12 images (1 variant from 3 parent and 3 hybrid categories, and phase-scrambled versions of these) was also used to familiarize participants with the rating task. Each batch was presented once to two participants. In total, 20 adults (7 female, age $M$ = 33.85, $SD$ = 12.53) rated images on the following scales: (1) a realistic image of a physical surface vs. an unrealistic pixelated pattern; (2) a flat, 2D texture vs. a 3D surface; (3) a soft surface vs. a hard surface; (4) a rough surface vs. a smooth surface; (5) a solid surface vs. a liquid or grainy material; (6) warm to touch vs. cold to touch; (7) wet to touch vs. dry to touch; (8) pleasant to touch vs. repulsive to touch; (9) overall easy vs. difficult to imagine how it would feel to touch. Each rating was made on a discrete scale from 1 to 6, with 1 corresponding to the first statement, and 6 corresponding to the second statement in the description of each scale. For example, if a participant perceived an image to be a highly realistic image of a physical surface, this would correspond to a rating of 1 on the first scale.

**Generative AI in the research process**. Stimulus images were generated using Midjourney (v5.2; Midjourney, Inc.), as described above. Experiment, data analysis, and data visualization code was developed with the assistance of Claude (Opus 5; Anthropic, PBC). All code was reviewed, tested, and validated by the authors, who verified that the reported results and figures were produced correctly and who take full responsibility for the analyses.
